# Internalized β2-Adrenergic Receptors Inhibit Subcellular Phospholipase C-Dependent Cardiac Hypertrophic Signaling

**DOI:** 10.1101/2023.06.07.544153

**Authors:** Wenhui Wei, Alan V. Smrcka

## Abstract

Chronically elevated neurohumoral drive, and particularly elevated adrenergic tone leading to β-adrenergic receptor (β-AR) overstimulation in cardiac myocytes, is a key mechanism involved in the progression of heart failure. β1-AR and β2-ARs are the two major subtypes of β-ARs present in the human heart, however, they elicit different or even opposite effects on cardiac function and hypertrophy. For example, chronic activation of β1ARs drives detrimental cardiac remodeling while β2AR signaling is protective. The underlying molecular mechanisms for cardiac protection through β2ARs remain unclear. Here we show that β2-AR protects against hypertrophy through inhibition of PLCε signaling at the Golgi apparatus. The mechanism for β2AR-mediated PLC inhibition requires internalization of β2AR, activation of Gi and Gβγ subunit signaling at endosomes and ERK activation. This pathway inhibits both angiotensin II and Golgi-β1-AR-mediated stimulation of phosphoinositide hydrolysis at the Golgi apparatus ultimately resulting in decreased PKD and HDAC5 phosphorylation and protection against cardiac hypertrophy. This reveals a mechanism for β2-AR antagonism of the PLCε pathway that may contribute to the known protective effects of β2-AR signaling on the development of heart failure.

---

Catecholamines of the sympathetic nervous system (SNS) regulate heart rate, contractility and vascular resistance through activation of adrenergic receptors. Prolonged elevation of circulating catecholamines, including epinephrine and norepinephrine, in response to cardiac injury, vascular disease, or stress is tied closely to the pathogenesis of heart failure (*1*). These hormones act in part through β-adrenergic receptors (βARs) on cardiac myocytes, where chronic activation drives maladaptive cardiac hypertrophy, apoptosis, and fibrosis, ultimately resulting in heart failure (*2*). βARs are G protein-coupled receptors (GPCRs) and consist of three subtypes, β1, β2 and β3. β1-adrenergic receptors (β1ARs) comprise of 80% of βARs in the healthy human hearts with the β2ARs comprising the remaining 20% (*2*). Although β1ARs and β2ARs respond to the same physiologic agents, share high sequence homology (*3*), and core signaling pathways (both couple to Gα_s_ and stimulate cAMP production), they have distinct or even opposite physiological and pathological roles in the heart (*4*). Specifically, chronic pathological stimulation by endogenous catecholamines leads to decreased β1AR levels and function in the heart, and mild overexpression of β1ARs results in cardiac hypertrophy and heart failure in mice (*5, 6*). In contrast, the β2AR population remains unchanged during heart failure and moderate levels of β2AR overexpression leads to positive inotropic effects (*7-10*). β2AR deletion in mice leads to exaggerated cardiac hypertrophy in a pressure overload-induced model, and β1AR deletion mice had a comparable responses to wild type, however, in β1AR/β2AR dual knockout mice, cardiac hypertrophy was abolished (*11, 12*). These observations suggest that coordinated signaling events downstream these receptors are critical in mediating cardiac hypertrophy and heart failure.

Phospholipase C (PLC) enzymes have important roles in the heart (*13-18*). PLCs mediate hydrolysis of phosphatidylinositol 4,5 biphosphate (PIP_2_) downstream of GPCR and receptor tyrosine kinase (RTKs) activation leading to production of inositol triphosphate (IP3) and diacylglycerol (DAG) (*19, 20*). Classically, PLCβ isoforms of the PLC family are stimulated downstream of GPCRs by Gq and Gβγ subunits from Gi (*21*). More recently, PLCε has also been shown to be downstream of multiple GPCR families and RTKs due to its ability to be directly activated by a diverse array of upstream regulators including multiple members of the family of small GTPases (Ras, Rho and Rap) and Gβγ subunits, (*22-25*).

A pathway that has been extensively studied in cardiac myocytes in our laboratory is regulation of PLCε by Rap after cAMP-dependent stimulation of the Rap GEF, Exchange Protein Activated by cAMP (EPAC) (*16-18, 26*). We previously identified an essential role of PLCε in regulating cardiac hypertrophy in an animal model of pressure-overload induced cardiac hypertrophy (*18*). In cardiac myocytes, PLCε scaffolds to muscle-specific A kinase anchoring protein (mAKAP) along with PKA, EPAC, PKD and other hypertrophic regulatory proteins at nuclear envelope (*27, 28*). Activation of PLCε at this location induces the hydrolysis of phosphatidylinositol-4-phosphate (PI4P) in the closely associated Golgi apparatus, generating inert inositol bisphosphate (IP2) and local DAG to drive the activation of PKD at nucleus and phosphorylation of HDAC leading to the hypertrophic gene expression (*18*).

While exploring mechanisms for cAMP-dependent stimulation of the Golgi Epac/PLCε pathway we found that stimulation of cell surface β1ARs with Iso was ineffective in cardiac myocytes (*29*). Rather, adrenergic stimulation of the Golgi Epac/PLCε pathway required a pre-existing pool of Golgi localized β1-ARs (*26*) which stimulate production of a pool of cAMP with privileged access to the Epac/PLCε module at the Golgi/NE interface. Access of Epi and NE to the intracellular β1-ARs requires an organic cation transporter subtype, OCT3 (*30, 31*) and blockade of OCT3 attenuated catecholamine-stimulated cardiomyocyte hypertrophy (*26*) and contractile responses (*32*). These studies clearly demonstrate a physiologically relevant roles of intracellular β1-ARs, however, how plasma membrane/sarcolemma β-AR signaling coordinates with intracellular β1-AR signaling remains poorly explored.

It is well established that intracellular cAMP pools generated at different subcellular compartments downstream of Gs coupled receptors lead to distinct phenotypic outcomes. In cardiac myocytes, β1-ARs but not β2-AR localize on the periphery of the nuclear envelope (*33, 34*) or Golgi (*26, 32, 35*) in addition to cell surface and T-tubule locations of both β1 and β2ARs. Activation or β2-ARs results in internalization into endosomes where they continue to signal, initiating a set of signaling and transcriptional control events distinct from those at the plasma membrane (*36*). Internalization of activated β2-ARs in cardiac myocytes has been documented (*37-39*) but functional roles for these internalized receptors have not been thoroughly examined in cardiomyocytes.

In this study, we demonstrate that activation of plasma membrane β2-ARs inhibits stimulation of the hypertrophic EPAC/PLCε pathway at the Golgi downstream of Golgi-β1-ARs and plasma membrane angiotensin II receptors in cardiac myocytes. We describe a mechanism where activated β2-ARs internalize from the plasma membrane, activate Gi, releasing Gβγ, subunits leading to ERK activation and inhibition of PI hydrolysis at Golgi apparatus. This prevents PLCε-mediated downstream hypertrophic signaling including nuclear PKD activation and HDAC phosphorylation. These data reveal a new mechanism that could underly the anti-hypertrophic versus hypertrophic signaling balance between β1 and β2-ARs and give insights into novel strategies for treatment of heart failure by deliberate inhibition of internal β1-ARs and combined with selective β2-AR activation for heart failure therapy.

## Results

### Activation of β2-ARs opposes Golgi-β1-AR-mediated PLCε activation at the Golgi apparatus

We previously demonstrated that the cell permeant agonist dobutamine (Dob) stimulates PLCε-dependent PI4P hydrolysis through activation Golgi β1ARs.(*26*). To assay Golgi PLCε activity we transduce cells with adenoviruses expressing GFP-FAPP(four phosphate adapter protein)-PH which binds selectively to PI4P and measure stimulus-dependent alterations in Golgi associated fluorescence using live-cell time lapse confocal fluorescence imaging in both neonatal rat ventricular myocytes (NRVMs) and adult ventricular myocytes (AVMs) (*18, 26*). Figure 1A shows co-localization of the PI4P-specific fluorescent probe with the Golgi marker GM130. Regions of interest corresponding to the Golgi apparatus in proximity to the nucleus are quantitated before and after agonist addition. As further evidence that Golgi PLCε activation by Dob requires Golgi localized β1ARs in AVMs we expressed Golgi-targeted-mApple-NB80 (enos-mApple-NB80) in AVMs to block downstream engagement of Gs by Golgi βARs, and monitored PI4P hydrolysis. Localization of enos-mApple-NB80 at the Golgi apparatus was confirmed by co-localization AVMs with the Golgi specific sensor FAPP (Fig S1A). Expression of enos-mApple-NB80 blocked Dob-stimulated PLCε activation while the control enos-mApple had no effect (Fig S1B). This recapitulates previous results demonstrating that Golgi βARs mediate activation of Golgi PLCε by β1AR agonists in AVMs.

**Figure 1.**
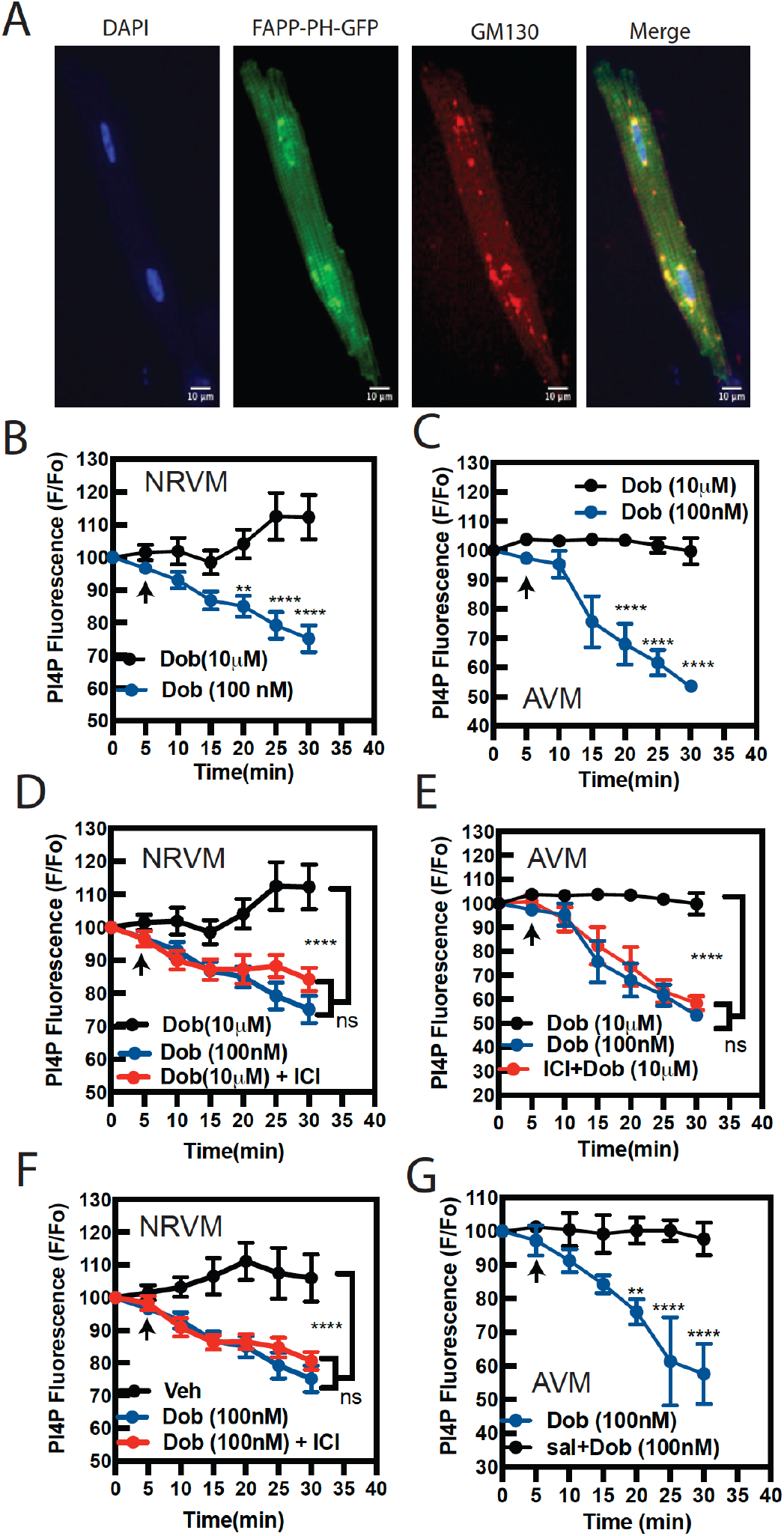
A pathway downstream of β2ARs suppresses β1AR stimulation of PLCε, at the Golgi. **A)** AVMs were transduced with adenoviruses expressing FAPP-PH-GFP for 18 hours prior to fixation and staining for GM130, a cis-Golgi marker. **B)** NRVMs were transduced with FAPP-PH-GFP and imaged by a live cell confocal microscopy. NRVMs were stimulated with dobutamine at either 10 *µ*M or 100 nM at the arrow and remaining fluorescence intensity at the Golgi apparatus was monitored over time. **C)** AVMs transduced with FAPP-PH-GFP were stimulated with dobutamine at either 10 *µ*M or 100 nM at the arrow indicated and Golgi associated fluorescence was monitored over time. NRVMs **(D and F)** and AVMs **(E)** were pretreated with a selective β2AR antagonist ICI-118,551 (50 nM) for 30 min before stimulation with dobutamine at the indicated concentrations and Golgi PI4P hydrolysis was measured. **G)** AVMs were pretreated with either Sal (100 nM) or vehicle control 30 min before imaging and dobutamine was added at the arrow. Images taken for PI4P hydrolysis were collected from n=3-5 cells from at least 3 independent preparations of AVMs and at least n=8 cells from 3 separate preparations of NRVMs. Data was analyzed with a two-way unpaired ANOVA with Sidak’s post-hoc test. *p<0.05; ** p<0.001; **** p<0.00001 using GraphPad Prism 9.

**Figure S1.**
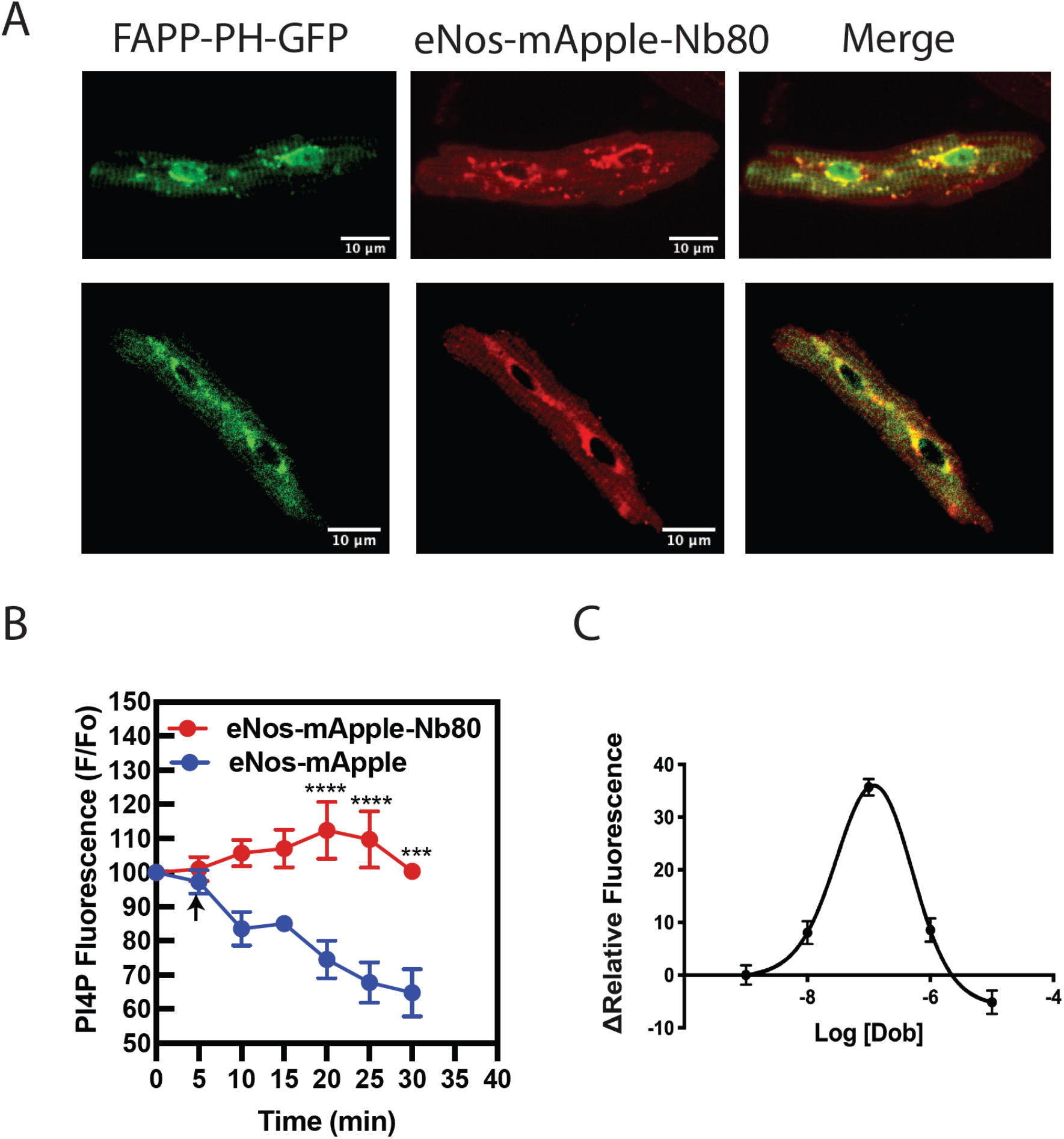
β1-ARs at Golgi apparatus are required for Dob-mediated PI4P hydrolysis in AVMs. **A)** Confocal images of AVMs co-expressing eNos-mApple-Nb80 (Golgi-targeted β-AR inhibitor), delivered in vivo using AAV9, and FAPP-GFP-PH delivered in vitro as described. **B)** AVMs expressing either eNos-mApple-Nb80 or eNos-mApple treated with Dob (addition indicated at arrow). Golgi associated FAPP-PH-GFP fluorescence was measured over time. Data was analyzed using an unpaired two-way ANOVA with Sidak’s post-hoc test. ***p<0.0001; ****p<0.00001. **C)** Concentration-dependence for dobutamine stimulated PI4P hydrolysis. NRVMs were transduced with FAPP-PH-GFP, the indicated concentrations of dobutamine were added and the change in Golgi associated PI4P fluorescence relative to time=0 was quantified after 25 min of dobutamine addition.

Unexpectedly, while analyzing the concentration-dependence for Dob-mediated Golgi PLCε activation, we found that 100 nM Dob was optimal, but 10 *µ*M Dob did not stimulate PI4P hydrolysis in NRVMs or AVMs (Fig 1B, C, S1C). Dob is relatively selective β1AR agonist but activates β2ARs at higher concentrations (pKd 5.6 vs 4.8, β1ARs vs β2ARs)(*40*). To assess whether the inhibitory effect on PI4P hydrolysis at higher concentrations of Dob was due to activation of β2ARs, we pretreated NRVMs or AVMs with the selective β2AR antagonist ICI-118,551 prior to Dob addition. Preincubation with ICI-118,551 uncovered stimulation of PI4P hydrolysis by 10 μM Dob comparable to that seen with 100 nM Dob in both NRVMs and AVMs (Fig. 1D, E). ICI-118,551 alone has no effects on 100 nM Dob-stimulated PI4P hydrolysis confirming that Dob-stimulated PI4P hydrolysis is β1AR-dependent (Fig. 1F). To extend this observation, we determined if salmeterol (Sal), a selective β2AR agonist, could inhibit PI4P hydrolysis stimulated by Dob. Preincubation with Sal (100 nM) inhibited PI4P hydrolysis stimulated by 100 nM Dob (Fig. 1G). Taken together, this data indicates that β2AR stimulation opposes PLCε activation downstream of Golgi β1ARs.

### β2AR-dependent inhibition of PI4P hydrolysis is at the level of PLCε, signaling

Previous studies from our laboratory demonstrated that Golgi β1AR-induced PI4P hydrolysis requires the EPAC activation downstream of βARs and upstream of PLCε, in NRVMs (*26*). To determine if β2ARs inhibit PI4P hydrolysis downstream of β1AR-dependent cAMP accumulation, we stimulated PLCε,-dependent PI4P hydrolysis using the EPAC-selective cAMP analog, 8-(4-chlorophenylthio)-2 ′ -O-methyl-cAMP-acetoxymethyl ester, (cpTOME-AM). Pretreatment with Sal prevented cpTOME-AM-stimulated Golgi PI4P hydrolysis indicating that β2ARs inhibit PLCε, activation at the level of EPAC or its downstream effectors, not at the level of cAMP generation by Golgi β1ARs (Fig 2A, B).

**Figure 2.**
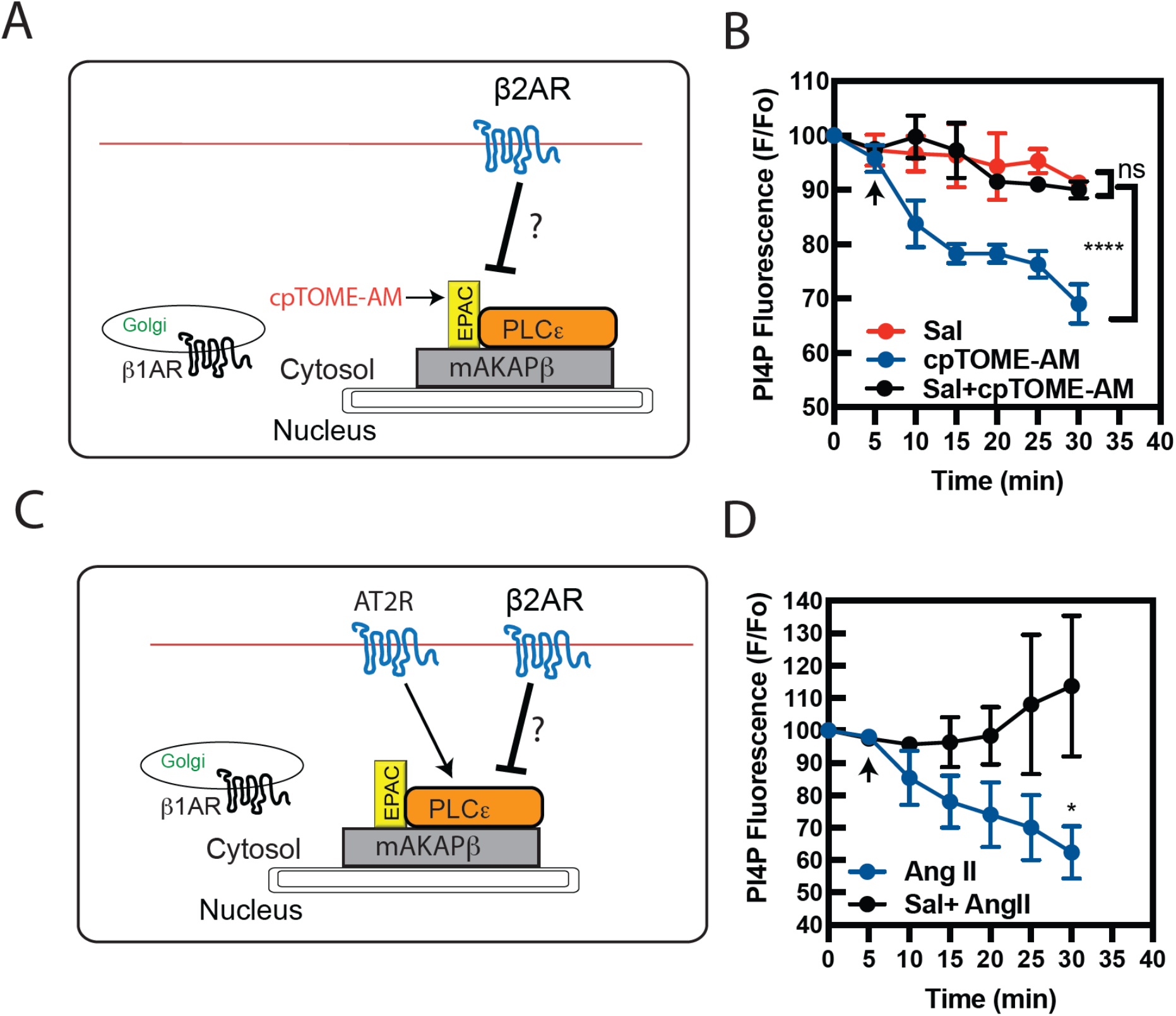
PLCε, is the likely target for β2AR-dependent inhibition of Golgi PI4P hydrolysis. **A)** Diagram of β2AR-dependent blockade of cpTOME-AM mediated activation of EPAC/PLCε,. **B)** AVMs were pretreated with or without Sal (100 nM) before stimulation with cpTOME-AM (10 *µ*M) and Golgi PI4P associated fluorescence was monitored with time. **C)** Diagram of β2AR-dependent blockade of AngII mediated activation of PLCε, at the Golgi apparatus. **D)** AVMs were pretreated with or without Sal (100 nM) before stimulation with Angiotensin II (1 *µ*M) and Golgi PI4P associated fluorescence was monitored. Images for PI4P hydrolysis were taken from n=3-8 cells from 3 independent preparations of AVMs. Data was analyzed with a two-way unpaired ANOVA with Sidak’s post-hoc test. *p<0.05; ****p<0.00001 using GraphPad Prism 9.

We previously demonstrated that a Gq coupled receptor (endothelin 1 A receptor) activates mAKAP associated PLCε, through a pathway independent from EPAC (*18*). We reasoned that if the inhibitory effect of β2AR activation was at the level of PLCε, beyond EPAC/Rap, activation of PLCε, by an alternative pathway would also be inhibited (Fig 2C). To test this AVMs were treated with Angiotensin II (AngII), which activates ATII receptors that couple to Gq and Gi/o but not Gs and is highly relevant to cardiac pathophysiology. AngII strongly stimulated PI4P hydrolysis which was blocked by Sal pretreatment (Fig 2D). This indicates that β2AR-dependent inhibition is at the level of PLCε,/mAKAP complex because two independent signaling pathways involving either Rap or signals downstream of Gq converge at the level of PLCε, activation at the Golgi.

### β2AR-dependent inhibition relies on Gi-Gβγ signaling

Activation of β2ARs stimulates cAMP production and PKA activation. We have reported that PKA activation counters the stimulation of Golgi PLCε, by EPAC (*29*). We tested whether PKA is involved in β2AR-dependent PLCε, inhibition (Fig. 3A). Preincubation AVMs with a PKA inhibitor, PKI, did not uncover PI4P hydrolysis upon stimulation with 10 μM Dob indicating that β2AR-mediated inhibition is PKA independent (Fig. 3B). β2ARs preferentially activate Gs, but also couple to Gi/o (*37, 41*). To test the role of Gi we treated AVMs with the Gi/o inhibitor, pertussis toxin (PTX). PTX treatment blocked Sal-mediated inhibition of PI4P hydrolysis induced by Dob (Fig. 3C). We then tested the role of Gβγ liberated from Gi/o activation downstream of β2ARs in the signaling events. Treatment with gallein, a Gβγ inhibitor (*42, 43*), prevented Sal-dependent inhibition of Dob-stimulated PLCε, activation (Fig. 3D).

**Figure 3.**
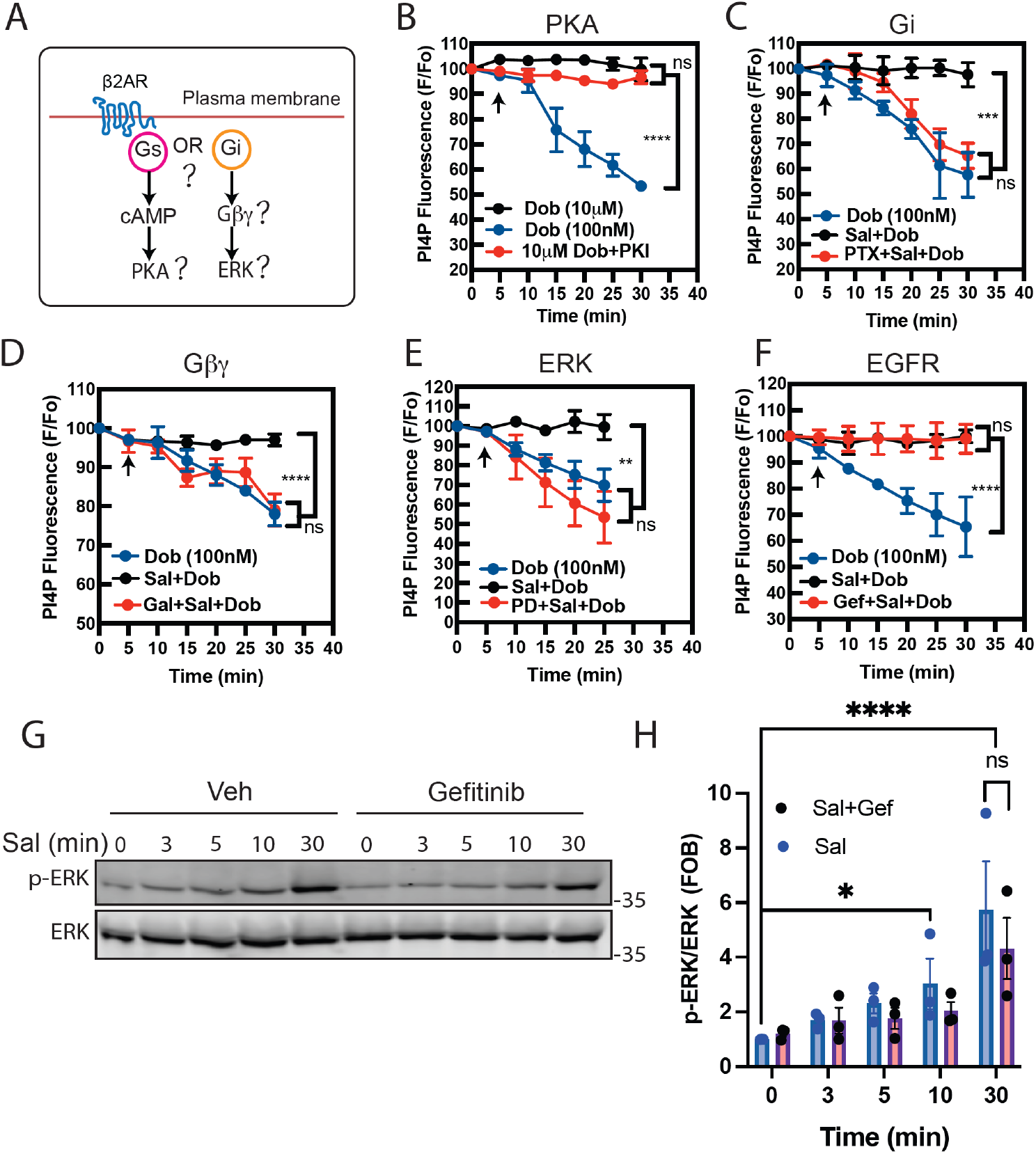
β2AR-Gi-Gβγ-ERK signaling axis counters activation of PLCε. **A)** Diagram of possible signaling pathways downstream of β2ARs. **B)** AVMs were pretreated with or without the PKA inhibitor, myrPKI (1 *µ*M) 30 min before stimulation with dobutamine at the indicated concentrations and Golgi PI4P hydrolysis was assessed. **C)** AVMs were pretreated with either PTX (100 ng/ml) or vehicle control overnight and followed by pretreatment with Sal 30 min before imaging and dobutamine was added at arrow. **D)** AVMs were pretreated with the Gβγ inhibitor, gallein (10 *µ*M) or vehicle control for 30 min prior to the pretreatment with Sal for 30 min and dobutamine was added at the arrow. **E)** AVMs were pretreated with the ERK inhibitor, PD0325901 (100 nM) for 30 min prior to the pretreatment with Sal for 30 min and dobutamine was added at the arrow. **F)** AVMs were pretreated with or without the EGF receptor inhibitor, gefitinib (10 *µ*M) for 2 hours before the pretreatment with Sal for 30 min and dobutamine was added at the arrow. **G-H)** Acutely isolated adult ventricular myocytes were pretreated or without gefitinib for 2 hours before the treatment with Sal for indicated times followed by western blotting for p-ERK and total ERK. Shown is a representative western blot from three independent preparations of AVMs with quantitation shown in H. Images for PI4P hydrolysis were taken from n=3-6 from 3 independent preparations of AVMs. Data was analyzed with a two-way unpaired ANOVA with Sidak’s post-hoc test. *p<0.05; **p<0.001; ***p<0.0001; ****p<0.00001 using GraphPad Prism 9.

### β2ARs inhibit PLCε, activation via ERK signaling

Given that β2ARs can signal through β-arrestin, a nexus for activation of protein kinases including ERK, we tested whether ERK activation downstream of β2ARs is involved. Pharmacological inhibition of ERK activity with PD0325901 abolished Sal-mediated inhibition of Golgi PI4P hydrolysis (Fig. 3E). Previous studies have reported that β1 and β2ARs can stimulate RTK (EGFR) transactivation, leading to ERK activation (*44-46*). This pathway requires the release of Gβγ subunits and is PTX sensitive. Additionally, βAR-mediated EGFR transactivation confers protective effects in isolated NRVMs (*47*). To assess the involvement of β2AR-EGFR transactivation, AVMs were pretreated with an EGFR inhibitor, gefitinib followed by preincubation with or without Sal followed by 100 nM Dob stimulation. EGFR inhibition did not alter the Sal-dependent inhibition of Dob-mediated PI4P hydrolysis (Fig. 3F). We also examined if β2AR-EGFR transactivation contributes to ERK activation downstream of β2ARs activation. AVMs were stimulated with Sal over a time course of 0-30 min with or without gefitinib preincubation and ERK phosphorylation was assessed by western blotting. Sal induced ERK activation weakly at 3-10 min, that appears to be inhibited by gefitinib. Sal treatment leads to statistically significant ERK activation at 30min and EGFR inhibition with Gefitinib had no statistically significant effect on this stimulation. (Fig. 3 G, H). These observations favor a model where β2ARs activate Gi and release Gβγ subunits, leading to EGFR-independent ERK activation that antagonizes Golgi PLCε. activation.

### Internalized β2ARs are required to activate ERK and inhibit PLCε. signaling

β2ARs undergo activation-dependent internalization which has been implicated in ERK activation (*48*). Inhibition of receptor internalization using a dynamin inhibitor, dyngo-4a markedly reduced the β2AR-mediated blockade of PLCε. activation (Fig. 4A) suggesting β2ARs mediate PLCε. inhibition from endosomes rather than the plasma membrane. Salmeterol is a partial β2AR agonist for both Gs and β-arrestin recruitment relative to Iso, yet does engage arrestin to some degree (*49, 50*). To determine if Sal causes β2AR internalization in cardiac myocytes, NRVMs were transfected with flag-β2ARs and internalization was monitored using fluorescent anti-flag M1 antibody and confocal microscopy. Stimulation of cells with Sal led to accumulation of intracellular β2AR associated fluorescence confirming that Sal does cause β2AR internalization in myocytes (Fig 4B). To determine if endocytic blockade affects β2AR stimulation of ERK phosphorylation, freshly isolated AVMs were pretreated with or without dyngo-4a and time-dependent ERK activation stimulated by Sal was measured by western blotting. Dyngo-4a significantly inhibited Sal-elicited ERK phosphorylation at 30 min consistent with a mechanism requiring receptor internalization (Fig. 4C, D). To explicitly examine the involvement of the endosomal Gβγ in the regulation of Golgi PLCε, activity, we selectively perturbed endosomal Gβγ using the C-terminus of G protein-coupled receptor kinase (GRK2ct), a highly selective Gβγ inhibitor, fused to a 2XFYVE domain sequence, highly specific to for binding to PI3P enriched endosomes (*51*), to generate an endosomal targeted GRK2ct (FYVE-mApple-GRK2ct) (Fig. 4E). Expression of FYVE-mApple-GRK2ct in AVMs led to a punctate expression pattern consistent with an endosomal location (Fig. 4F). In AVMs co-expressing FAPP-PH-GFP and FYVE-mApple-GRK2ct, Sal was unable block Dob-induced PI4P hydrolysis (Fig. 4G) indicating that the inhibitory signaling downstream of β2ARs is indeed driven by endosomal Gβγ.

**Figure 4.**
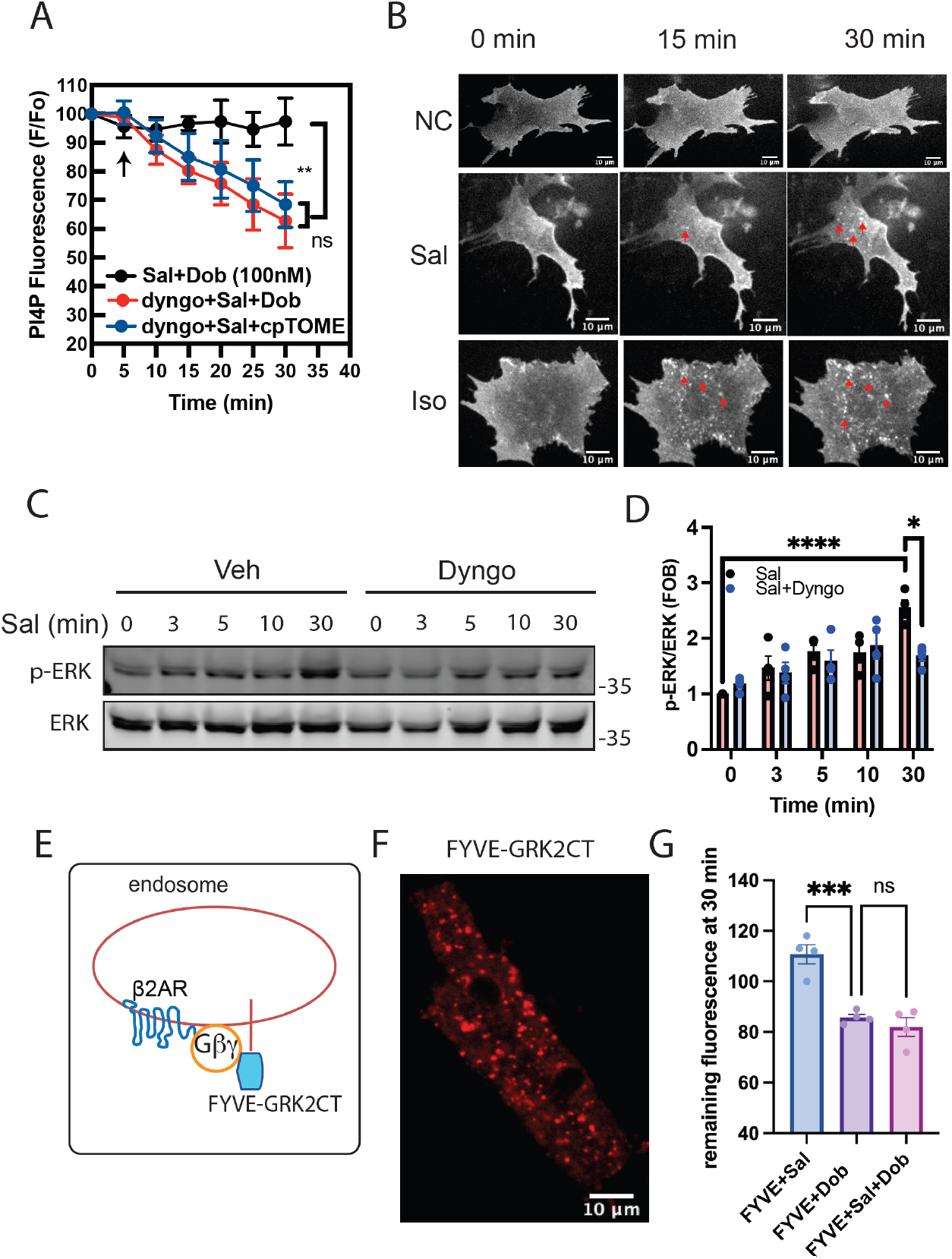
β2AR-dependent blockade of PLCε, activation relies on endosomal Gβγ. **A)** AVMs were pretreated with the dynamin inhibitor Dyngo-4a (40 *µ*M) for 30 min prior to pretreatment with Sal for 30 min and dobutamine was added at the arrow. Data was analyzed with a two-way unpaired ANOVA with Sidak’s post-hoc test **p<.001, ns=not significant. **B)** Representative images showing Sal or Iso mediated β2-AR internalization. NRVMs were transfected with Flag-β2ARs for 24 hours. Cells were then labeled with M1-Flag-488 for 10min at 37°C and then treated with negative control, Sal (100 nM), or Iso (10 *µ*M) at 37°C. Images were acquired at the indicated times by confocal microscopy. **C-D)** Acutely isolated adult ventricular myocytes were pretreated in the presence or absence of Dyngo-4A before the treatment with Sal for indicated times followed by western blotting for p-ERK and total ERK. Shown is a representative western blot from four independent preparations of AVMs with quantitation shown in D. Data was analyzed using an unpaired two-way ANOVA with Sidak’s post-hoc test. *p<0.05; ****p<0.00001. **E)** Diagram of blockade of Gβγ signaling at endosomal membranes. **F)** Representative image of AVMs expressing FYVE-GRK2ct. **G)** AVMs were transduced with adenoviruses expressing FYVE-mApple-GRK2ct and FAPP-PH-GFP for 18 hours before imaging. AVMs were stimulated with Dobutamine alone or in the presence of Sal. Golgi associated PI4P fluorescence intensities at 0 and 30 min were measured. Images for PI4P hydrolysis were taken from n=3-9 cells from at least 3 independent preparations of AVMs. Data was analyzed with a one-way ANOVA with Sidak’s post-hoc test. *** p<0.0001 using GraphPad Prism 9.

### Activation of β2ARs inhibits nuclear PKD activation

A signaling event directly downstream of perinuclear PLCε, is the activation of nuclear PKD (*18*). PKD is directly activated by DAG, the principal active product of PLCε, activity acting on Golgi PI4P. If Golgi PLCε, is inhibited downstream of Sal, we would predict that β2AR activation would inhibit nuclear PKD activation. To determine if activation of β2ARs reduces agonist-dependent PKD activation, NRVMs were pretreated with Sal for 30 min before addition of AngII for 30 min and analysis of PKD phosphorylation by western blotting. Sal treatment eliminated AngII mediated PKD activation (Fig. 5A, B). To more specifically examine nuclear PKD activation downstream of β2AR and Ang II activation, AVMs were transduced with an adenovirus expressing a nuclear-localized fluorescence resonance energy transfer (FRET) PKD activation reporter, nDKAR. (*52*) (Fig. 5C). AVMs were pretreated with either vehicle or Sal for 30 min and AngII-mediated changes in nuclear FRET was measured in the nucleus at 0 and 30 min. AngII caused a significant nuclear PKD activation which was suppressed by β2AR activation (Fig. 5D).

**Figure 5.**
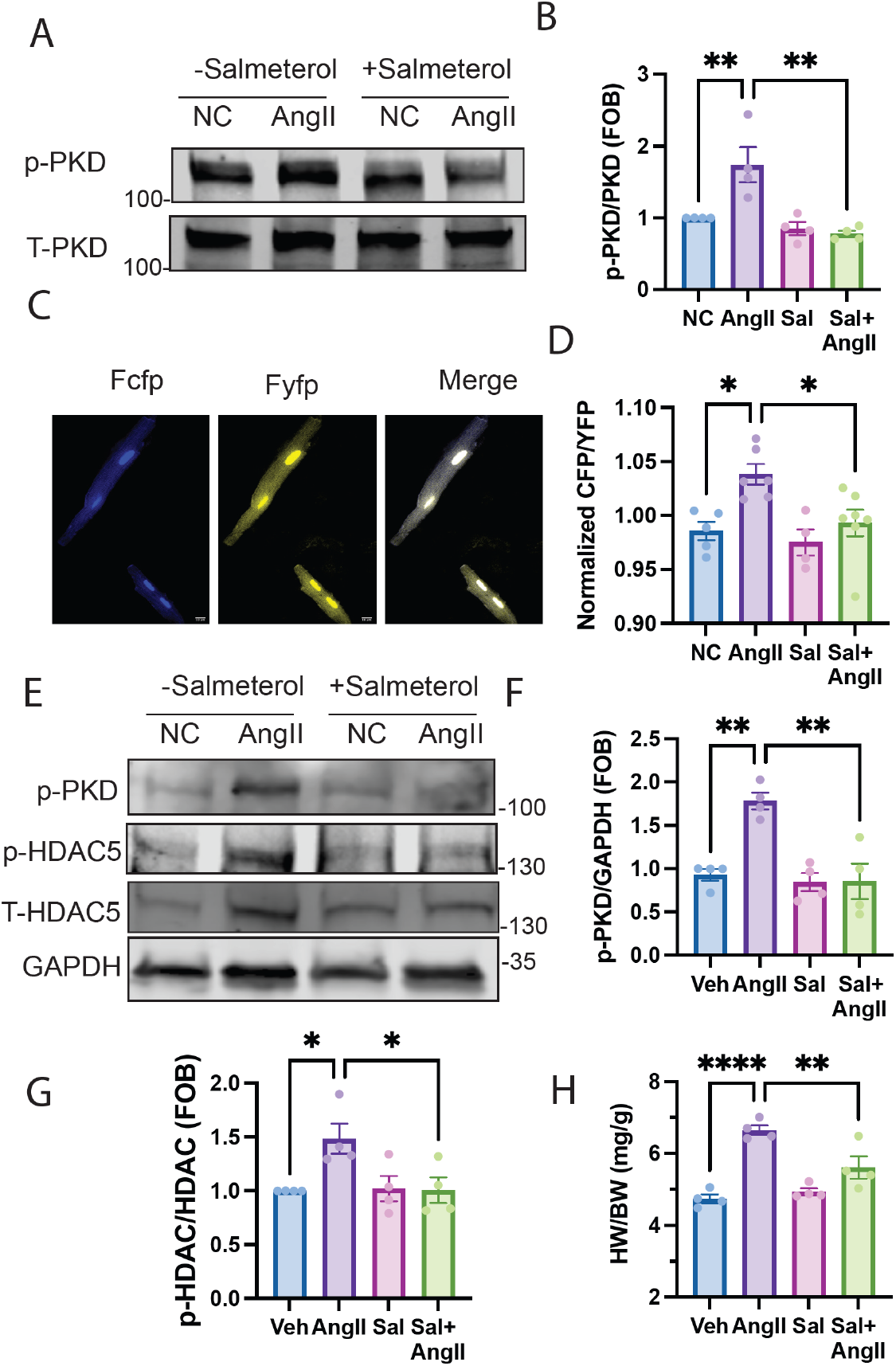
Nuclear PKD activation downstream of PLCε, is suppressed by β2AR activation. **A-B)** NRVMs were pretreated with either Sal or vehicle before addition of AngII (1 *µ*M) for 30 min and followed by western blotting to for p-PKD and total PKD. Shown is a representative western blot from four separate preparations of NRVMs. Quantitation is shown in B. **C)** Representative images of AVMs expressing nuclear-DKAR. **D)** The nuclear region of AVMs expressing nDKAR was selected and the CFP/YFP ratio was measured before and after addition of AngII (1 *µ*M) for 30 min addition in the presence with or without pretreatment with Sal for 30 min. **E)** Heart lysates from WT mice infused with either saline (Veh) or salmeterol (Sal) (25 *µ*g/kg/day) together with or without AngII (1.5 mg/kg/day) for 14 days. Western blotting was performed to determine the level of p-PKD, p-HDAC, total HDAC and GAPDH. Shown is a representative western blot from 4 mice each group. **F)** Quantitation of PKD phosphorylation from E. **G)** Quantitation of HDAC phosphorylation from E. H) Heart weight/body weight (HW/BW) ratios from AngII and AngII+Sal treated mice. N=4 animals each condition. Data was analyzed with a one-way ANOVA with Sidak’s post-hoc test. *p<0.05; **p<0.001; **** p<0.00001 using GraphPad Prism 9.

We had previously shown that cardiac specific deletion of PLCε, in mice eliminates PKD activation in a TAC model suggesting that PKD phosphorylation can be used as a proxy for PLCε, activity in vivo. To extend our observations to mice, we examined PKD phosphorylation in response to the chronic AngII infusion together with Sal by implanting AngII +/- Sal subcutaneous osmotic minipumps into mice for 14 days. Chronic stimulation with Sal significantly attenuated AngII-mediated PKD activation in mouse hearts (Fig. 5E and F). Similarly, HDAC phosphorylation was blunted in Sal treated group (Fig. 5E and G). PKD phosphorylation of HDAC is partially responsible mediating MEF-dependent hypertrophic gene transcription (*53*). Salmeterol also significantly inhibited AngII-driven cardiac hypertrophy as measured as a reduction in heart size (HW/BW) compared to animals treated with AngII alone (Fig. 5H). These observations together suggest β2ARs inhibit development of hypertrophy in mice by preventing detrimental AngII-mediated PLCε, signaling in cardiomyocytes. However, we cannot exclude the possibility that Sal mediated protection in mice involves inhibition of AngII-mediated hypertension.

### Dob-induced NRVM hypertrophy is effectively inhibited by β2ARs activation

In our previous studies, we showed that Golgi resident β1AR signaling to PLCε, is required for catecholamine-mediated cardiomyocyte hypertrophy in NRVMs (*26*). NRVMs were treated with Dob (100nM) with or without Sal for 48 hours and cell area and expression of the hypertrophic marker, atrial natriuretic factor (ANF), were measured. Dob stimulated a significant increase in cell area (Fig 6A, B) and ANF expression (Fig 6C, D). Co-treatment with Sal blunted Dob-induced hypertrophy assessed by these two measurements (Fig 6A, B, C, D), demonstrating that β2AR signaling is protective against cardiomyocyte hypertrophy consistent with the results from mice.

**Figure 6.**
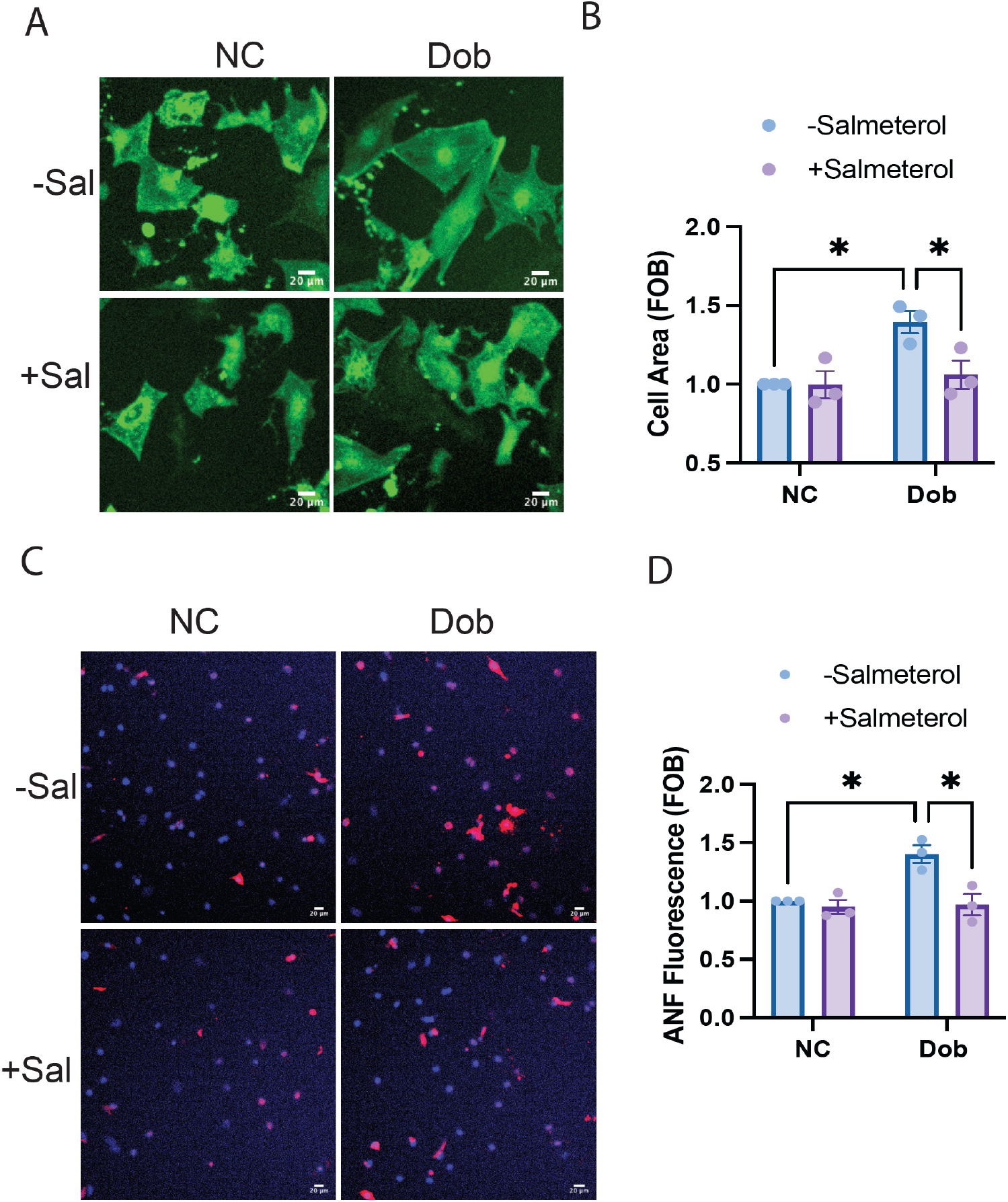
Activation of by β2ARs inhibits dobutamine induced cardiomyocyte hypertrophic growth. **(A-B)** NRVMs were stimulated with dobutamine for 48 hours in the presence of Sal or vehicle control and followed by fixation and staining for α-actinin to identify cardiomyocytes and quantitated for cell area by image J. **(C-D)** NRVMs were stimulated with dobutamine for 48 hours in the presence of Sal or vehicle control and followed by fixation, cells were stained for ANF expression and with DAPI to identify nuclei. The fluorescence intensity of ANF rings was quantified by image J. Data was quantified from at least n=350 cells from 3 separate preparations of NRVMs. Data was analyzed with a one-way ANOVA with Tukey’s post-hoc test. *p<0.05; *p<0.001 using GraphPad Prism 9.

## Discussion

Our previous work led to development of a model where stimulation of Golgi β1ARs by the membrane permeant agonist Dob produces a local pool of cAMP with privileged access to the EPAC/PLCε/mAKAP complex, generating DAG derived from PI4P depletion, activating PKD and mediating cardiac hypertrophy (*18, 26*). Unexpectedly, while low concentrations of Dob induced robust PI4P hydrolysis, saturating concentrations of Dob did not. Here we demonstrate that β2AR activation opposes activation of Golgi PLCε-PKD pathway by two clinically relevant hypertrophic stimuli which could play an essential role in the ability of β2ARs to limit cardiac hypertrophy. In cardiac myocytes we show that the mechanism involves agonist-driven internalization of β2ARs where they couple to ERK activation via Gi-Gβγ at the endosome which in turn inhibits prohypertrophic PLCε signaling at the Golgi. Heart failure is often associated with up-regulation of Gi and enhanced β2AR-Gi signaling (*54, 55*) which may be a compensatory mechanism to overcome the detrimental effects induced by circulating mediators and sympathetic neurotransmitters, many of which are likely to signal through PLCε.

### Compartment specific ERK signaling in the heart

Extensive studies suggest ERK is a key player in regulating cardiomyopathy. ERK signaling can produce both the beneficial and deleterious effects depending on the context. Transgenic mice overexpressing activated ERK are reported to show maladaptive (*56*) or adaptive (*57*) hypertrophic responses, while other studies utilizing mice with ERK deletion mice suggest ERK1/2 signaling may be dispensable in pathologic cardiac hypertrophy (*58, 59*). In addition, an in vitro study indicates that ERK1/2 activation is required for the ANF mediated antihypertrophic response (*60*). It is now becoming clear that the signaling cascades and outcomes downstream of ERK largely depend on the subcellular locations of phosphorylated ERK1/2 (*61*). Therefore, a more sophisticated dissection of ERK signaling mechanisms is required.

ERK is activated downstream of multiple GPCRs which are themselves spatially compartmentalized (*62*). GPCRs inside the cell activate G proteins and produce local cAMP accumulation to initiate distinct subcellular signaling events (*63, 64*). G proteins including Gs, Gq, Gi and Gβγ subunit or β-arrestin-mediated signaling pathways activate ERK cascades. Different ligands cause different subcellular destinations of activated ERK via their preference to shift GPCR coupling to G-proteins or β-arrestin (*65, 66*). Recently, it has been shown that after ligand-stimulated β2ARs endocytosis, endosomal ERK signaling is activated by an endosome-localized active Gα_s_ to subsequently stimulate nuclear ERK activity to control gene expression in HEK cells (*48*). Our studies support the idea that in AVMs, β2AR-induced accumulated ERK signaling requires receptor endocytosis and endosomal Gβγ released from Gi. This endosome-receptor initiated ERK signaling serves as a repressor for PLCε which has been implicated in mediating cardiac hypertrophy. This study provides a functional role for this endosomal β2AR-ERK signaling axis in preventing the development of cardiac hypertrophy.

### PKD signaling in cardiac remodeling

Cardiac-specific deletion of PKD improves cardiac function in response to pressure overload or angiotensin II signaling (*67*), making it a promising therapeutic target to treat heart failure resulting from maladaptive cardiac hypertrophic signaling. We previously demonstrated that DAG generated from PI hydrolysis by perinuclear scaffolded PLCε activates PKD in close proximity to the nucleus, where it phosphorylates HDAC to regulate hypertrophic gene expression (*18*). Our data suggest that acute activation of β2ARs can protect against pathological cardiac hypertrophy through inhibition of phosphorylation of nuclear PKD activation and perhaps other kinases such as CamKII. Hence, currently clinically effective β2AR agonists can serve as potent PKD repressors. In addition to the role of PKD in cardiac hypertrophy and fibrosis, it is also a critical signaling molecule in cancer-associated functions such as migration, cell proliferation, or survival (*68*). Thus our observations could extend to other pathophysiological systems.

Molecular mechanisms underlying the cardioprotective effects of β2AR signaling have been investigated previously. Gi-dependent-PI3K-AKT signaling downstream of β2-ARs contributes to its cardioprotective effects through prevention of apoptosis (*69*). In addition, β2-ARs can mediate EGFR and PDGFR transactivation, promoting cardiomyocyte survival (*70*). These studies focused on cardiomyocyte apoptosis. We propose that inhibition of hypertrophic PLCε signaling is an additional mechanism for β2AR dependent cardiac protection against detrimental cardiac remodeling.

### Combined therapy with a β2AR agonist and a selective β1AR blocker

β-blockers (carvedilol, bisoprolol, and metoprolol) are clinically used to treat patients with heart failure by blocking detrimental β1AR-G protein signaling, although their complete mechanism of action is not fully understood. More recently, it has been suggested that effective β-blockers tend to be hydrophobic which allows them to access the internal pools of β1ARs (*26, 35*). As a non-selective β-blocker, carvedilol has been to shown to produce superior clinical outcomes relative to other β-blockers (*71*), though the mechanism is unclear. Carvedilol has been suggested to activate β2ARs and ERK signaling (*72*). It has been proposed that carvedilol might have dual therapeutic benefits in heart failure by preventing the deleterious signaling from β1ARs and promoting the beneficial effects of β2ARs (*4*). This therapeutic concept combining β1AR blockade with a β2AR agonist has been previously proposed and investigated in a coronary ligation model mice and rats where the combination treatment was more effective at preserving cardiac function that β1 blockade alone (*69, 73, 74*). Our data indicate that in addition to effects on vascular tone selective β2ARs ligands can act directly through cardiac myocytes to improve cardiac function by preventing the deleterious PLCε stimulated by chronically elevated catecholamines and angiotensin.

### Isoproterenol paradox

Isoproterenol (Iso), a non-selective and relatively membrane impermeant β1AR and β2AR agonist, does not induce acute PLCε activation (*29*), but with chronic exposure, it causes cardiac hypertrophy in vitro and vivo. One possible explanation for this discrepancy could be with acute Iso treatment, β2AR activation counteracts signaling from β1ARs or AngII. However, with chronic exposure to strong stimulation by Iso β2ARs, while not downregulated, may undergo desensitization leading to the loss of their protective signaling. Alternatively, Iso may depend on a different plasma membrane mediated hypertrophic pathway. These speculations need to be experimentally verified.

## Materials and Methods

### Materials

Antibodies: mouse anti-α-actinin (Sigma, Cat# A7811), rabbit anti-ANF (Millipore, Cat# AB-5490), mouse anti-GM130 (BD Transduction Laboratories, Cat# 610822), mouse M1 anti-Flag (Millipore Sigma, Cat# F3040) labelled with Alexa Fluor 488 using Thermofisher labeling kit A20181 (gift from Dr. Manojkumar Puthenveedu), rabbit anti-p-PKD (Cell Signaling Technology, Cat# 2051), rabbit anti-t-PKD (Cell Signaling Technology, Cat# 2054), rabbit anti-p-ERK (Cell Signaling Technology, Cat# 4370S), mouse anti-t-ERK (Cell Signaling Technology, Cat# 4696S), rabbit anti-p-HDAC (Cell Signaling Technology Cat# 3433), mouse anti-t-HDAC (Santa Cruz Biotechnology Cat# sc-133225), mouse anti-GAPDH (Invitrogen, Cat# MA5-15738), goat anti-rabbit DyLight 800 (Invitrogen, Cat# SA535571), goat anti-mouse IRDye 680RD (LICOR, Cat# 926-68070), goat anti-mouse 568 secondary antibody (Invitrogen, Cat# A-11031), goat anti-rabbit 568 secondary antibody (Invitrogen, Cat# A-11011), goat anti mouse 488 secondary antibody (Invitrogen, Cat# A-11029).

Compounds: Cytosine Arabinoside (AraC, Sigma); Collagenase Type 2 (Worthington, Cat# LS004176); Gelatin (Sigma), Gallein (Sigma, 371708), Isoproterenol (Sigma, 1351005), Dobutamine (Tocris, Cat# 0515); ICI 118,551 (Tocris, Cat#0821), butanedione-monoxime (BDM) (Sigma, Cat# 112135), Dyngo (Abcam, Cat# Ab120689), pertussis toxin (Sigma, Cat# P7208-50UG), gefitinib (LC Laboratories, Cat# G-4408), myristoylated PKA inhibitor (Sigma, Cat# 476485), salmeterol xinafoate (Tocris, Cat# 47-1210), angiotensin II (Sigma, Cat# A9525-50mg), PD0325901 (Sigma, Cat# PZ0162-5mg), cptome-AM (Fisher scientific, Cat# 4853100U)

Plasmids and adenoviral constructs: FAPP-GFP-PH (*18*), GFP-P4M-SidMx2 (Addgene plasmid #51472), fyve mApple GRK2ct (created by fusing 2XFYVE domains (*75*), mApple, and GRK2CT (GRK2 (495-689)), made by twist bioscience), eNos-mApple-Nb80 (created by fusing the first 33 amino acids of enos (*76*), mApple and Nanobody 80, (made by twist bioscience), nDKAR (*18, 52*)

### Methods

#### Isolation and transduction of neonatal rat ventricular myocytes (NRVMs) and mouse adult cardiac myocytes (AVMs)

Neonatal rat ventricular myocytes were isolated from 2 to 4 day-old Sprague-Dawley rats as described previously (*27*). Briefly, ventricles were separated from the hearts and minced well before adding digestion buffer (Collagenase type II in Hanks buffered saline solution without calcium). Following digestion, supernatant was collected, and cells were centrifuged at 1200 rpm for 2 min. Fibroblasts were removed by pre-plating cells onto tissue culture plastic for one hour at 37°C. Purified NRVMs in the supernatant were transferred onto gelatin-coated glass-bottom dishes or 6 well plates. NRVMs were cultured in DMEM supplemented with 10% FBS, 100 U/mL penicillin, 100 ug/mL streptomycin, 2 μg/mL vitamin B12, and 10 μM cytosine arabinoside. 24 hours later, media were changed to 1% FBS or serum free media. For adenovirus transduction, 50 MOI of adenovirus was incubated overnight.

Isolation of adult ventricular myocytes (from 2-5 month-old wild type C57BL/6 mice) was performed following the protocol from (*77*). Mice were anesthetized with ketamine (100 mg/kg body weight) and xylazine (5 mg/kg body weight). Hearts were then removed, cannulated, and perfused with perfusion buffer (10 mM HEPES, 0.6 mM Na_2_PO_4_, 113 mM NaCl, 4.7 mM KCl, 12 mM NaCO3, 0.6 mM KH2PO4, 1.2 mM MgSO4, 10 mM KHCO3, 30 mM Taurine, 10 mM BDM, 5.5 mM Glucose) via the aorta. Subsequently, digestion buffer (collagenase type II (773.48 U/ml), trypsin (0.14 mg/ml), and calcium chloride (12.5*µ*M) in perfusion buffer) was perfused via the aorta. Following digestion, hearts were minced in stopping buffer (10% FBS and 12.5 *µ*M calcium chloride in perfusion buffer), debris was allowed to settle, the supernatant was collected, and calcium was added back to a final 1 mM concentration. Cells were then centrifuged at 18xg for 3 min before plating onto laminin-coated 20 mm glass bottom dishes for confocal microscopy imaging. AVMs were maintained in minimum essential medium (MEM) supplemented with 0.35 g/L sodium bicarbonate, 100 U/mL penicillin, 100 U/mL streptomycin, and 10 mM BDM. To transduce AVMs with adenovirus, 100 MOI of indicated adenoviruses were added to cells in BDM-free media for 4 hours. Following this, the virus was removed, and the culture media was supplemented with BDM. After 18 hours, cells were imaged by confocal microscopy.

#### Imaging for analysis of Golgi PI4P hydrolysis

NRVMs or AVMs were imaged at 37°C and 5% CO_2_ in a stage top incubator 18-24 hours after transduction with indicated adenoviruses in serum free culture media. Live cell imaging was performed on a Leica DMi8 microscope in spinning disc confocal mode (Crest Optics X-light V2) with 40X 1.4-NA (numerical aperture) oil immersion lens. CFP, GFP, and RFP were excited 440, 488 and 555 nm respectively with an 89-North LDI light source and emission was monitored and imaged on a backlit CMOS Photometrics Prime 95B camera.

Measurement of PI4P hydrolysis was previously described (*15, 18, 26, 29*). NRVMs or AVMs were transduced with adenovirus expressing FAPP-PH-GFP. Transduced cells were identified and imaged on a LEICA DMi8 with 40X oil lens in confocal mode. Timelapse videos were acquired with 10 ms exposure times at 5 min intervals to minimize phototoxicity and photobleaching. Fluorescence intensity analysis was performed by subtracting background cytosolic fluorescence from the fluorescence intensity corresponding to Golgi using Image J to circle the regions of interest corresponding to Golgi surrounding the nucleus in both AVMs and NRVMs. Agonists were added after 5 min baseline imaging. Data were normalized to the initial fluorescence intensity prior to addition of agonists. For PI4P measurement in FYVE-mApple-GRK2CT expressing AVMs, cells were excited with GFP and RFP to acquire images before and after 30 min of dobutamine addition with or without salmeterol preincubation. Data is presented as percentage of fluorescence remaining after dobutamine addition at 30 min. GFP: excitation: 488nm, emission: 510 nm. RFP: excitation: 555 nm, emission: 583 nm.

#### Measurement of myocyte cell area and ANF induction

NRVMs were plated onto gelatin-coated 6-well plates. Cells were then serum starved for 24 h before stimulation with dobutamine (100 nM) for 48 h co-incubated with or without salmeterol (100 nM). Cells were then fixed with 4% paraformaldehyde for 15min at room temperature and then incubated with 10% goat serum in 0.1% Triton X100 (PBST). α-actinin (1:100) or ANF antibodies (1:1000) were incubated overnight in 2% goat serum in PBST at 4°C. The following day, after three washes with PBST, secondary antibodies (goat anti-rabbit 568 secondary antibody or goat anti mouse 488 secondary antibody) were incubated for 1 h at room temperature at a dilution of 1:1000 in 2% goat serum in PBST. Fluorescence images was captured at 10X magnification. The area of the stained myocytes and ANF staining surrounding the nucleus were quantified using Image J software.

#### AAV9 mediated gene delivery

AAV9-eNos-mApple or AAV9-eNos-mApple-Nb80 were generated by University of Michigan Viral Vector Core. AAV9-eNos-mApple or AAV9-eNos-mApple-Nb80 were injected into the mediastinum of 7 day-old C57Bl/6 wild-type mice at a dose of 10^12^ viral genomes. Adult myocytes were isolated at 8 weeks of age and expression and localization were confirmed by confocal microscopy.

#### AngII mini pump

Angiotensin II was dissolved in saline with or without salmeterol xinafoate and were filled into osmotic mini-pumps (Alzet, model 1002). Pre-filled pumps were incubated in sterile saline at 37°C overnight. 8-10 week old male C57BL/6 mice were subcutaneously implanted with the mini-pumps to deliver angiotensin at the rate of 1.5 mg/kg/day and salmeterol at the rate of 25 *µ*g/kg/day for 14 days. Mice were then euthanized and heart weight to body weight ratio was determined. Heart lysates were analyzed by western blotting as indicated.

#### Western blotting

NRVMs or AVMs were lysed in 1X Laemmli sample buffer, boiled, and loaded onto 4-20% gradient mini-PROTEAN TGX gels (4561094, Bio-Rad). Proteins were then transferred to nitrocellulose membranes overnight at 25 mA or 2 h at 300 mA. Membranes were blocked with 4% bovine serum albumin (Fisher BP1600) in TBST (0.1% Tween-20 in Tris-buffered saline) for 2 hours. Primary antibodies were incubated overnight at 4°C followed by 3X washes with TBST. Secondary antibodies were incubated for 1 h at room temperature followed by 3X washes with TBST. Western blots were imaged and quantified with a LI-COR Odyssey imaging system.

#### Nuclear PKD FRET analysis

AVMs were transduced with adenovirus expressing nDKAR at 100 MOI. The following day, cells were imaged with LEICA DMi8 microscope at 37C in confocal mode with 40X oil lens. Transduced AVMs were identified with CFP channel and then stimulated with indicated agonists for 30min. FRET was measured as the ratio of YFP emission to CFP emission after CFP excitation. CFP, excitation: 440nm; emission: 480nm. YFP emission: 535nm. Due to the sensitivity of the myocytes to CFP excitation, data was collected at 0 and 30 min rather than a full time course.

#### Internalization of β2-adrenergic receptors

NRVMs were plated onto glass bottom dishes. After 24 h, cells were transfected with 500 ng Flag-β2-ARs using lipofectamine 3000 for 16 h. Cells were then incubated with M1-Flag-488 antibody for 10 min at 37°C. Labeled cells were identified in the GFP channel (excitation: 488nm, emission: 510 nm) and then stimulated with salmeterol for 30 min at 37°C. NRVMs were imaged in the GFP channel with a LEICA DMi8 microscope in confocal mode with 40X oil lens.

#### Statistics

Data were presented as mean +/- SEM from independent preparations of cells. Data were analyzed with a two way ANOVA with sidak’s post-hoc test or a one way ANOVA with Tukey’s multiple comparison test.

## Acknowledgements

We would like to thank the Brody lab especially Kumar Subramani, James Teuber, and Dr. Kobina Essandoh for sharing NRVMs and reagents. We would like to thank Dr. Manojkumar Puthenveedu lab especially Ian Chronis and Hao Chen for sharing reagents. This work is supported by grant R35 GM127303-01 (A.V.S.).

## References

1. M. R. Bristow, beta-adrenergic receptor blockade in chronic heart failure. Circulation 101, 558–569 (2000); published online EpubFeb 8 (10.1161/01.cir.101.5.558).

2. H. A. Rockman, W. J. Koch, R. J. Lefkowitz, Seven-transmembrane-spanning receptors and heart function. Nature 415, 206–212 (2002); published online EpubJan 10 (10.1038/415206a).

3. T. Frielle, S. Collins, K. W. Daniel, M. G. Caron, R. J. Lefkowitz, B. K. Kobilka, Cloning of the cDNA for the human beta 1-adrenergic receptor. Proc Natl Acad Sci U S A 84, 7920–7924 (1987); published online EpubNov (10.1073/pnas.84.22.7920).

4. A. Y. Woo, R. P. Xiao, beta-Adrenergic receptor subtype signaling in heart: from bench to bedside. Acta Pharmacol Sin 33, 335–341 (2012); published online EpubMar (10.1038/aps.2011.201).

5. J. D. Bisognano, H. D. Weinberger, T. J. Bohlmeyer, A. Pende, M. V. Raynolds, A. Sastravaha, R. Roden, K. Asano, B. C. Blaxall, S. C. Wu, C. Communal, K. Singh, W. Colucci, M. R. Bristow, D. J. Port, Myocardial-directed overexpression of the human beta(1)-adrenergic receptor in transgenic mice. J Mol Cell Cardiol 32, 817–830 (2000); published online EpubMay (10.1006/jmcc.2000.1123).

6. S. Engelhardt, L. Hein, F. Wiesmann, M. J. Lohse, Progressive hypertrophy and heart failure in beta1-adrenergic receptor transgenic mice. Proc Natl Acad Sci U S A 96, 7059–7064 (1999); published online EpubJun 8 (10.1073/pnas.96.12.7059).

7. C. A. Milano, L. F. Allen, H. A. Rockman, P. C. Dolber, T. R. McMinn, K. R. Chien, T. D. Johnson, R. A. Bond, R. J. Lefkowitz, Enhanced myocardial function in transgenic mice overexpressing the beta 2-adrenergic receptor. Science 264, 582–586 (1994); published online EpubApr 22 (10.1126/science.8160017).

8. G. W. Dorn, 2nd, N. M. Tepe, J. N. Lorenz, W. J. Koch, S. B. Liggett, Low- and high-level transgenic expression of beta2-adrenergic receptors differentially affect cardiac hypertrophy and function in Galphaq-overexpressing mice. Proc Natl Acad Sci U S A 96, 6400-6405 (1999); published online EpubMay 25 (10.1073/pnas.96.11.6400).

9. H. T. Tevaearai, A. D. Eckhart, G. B. Walton, J. R. Keys, K. Wilson, W. J. Koch, Myocardial gene transfer and overexpression of beta2-adrenergic receptors potentiates the functional recovery of unloaded failing hearts. Circulation 106, 124–129 (2002); published online EpubJul 2 (10.1161/01.cir.0000020220.79105.fd).

10. G. Rengo, C. Zincarelli, G. D. Femminella, D. Liccardo, G. Pagano, C. de Lucia, G. G. Altobelli, V. Cimini, D. Ruggiero, P. Perrone-Filardi, E. Gao, N. Ferrara, A. Lymperopoulos, W. J. Koch, D. Leosco, Myocardial beta(2) -adrenoceptor gene delivery promotes coordinated cardiac adaptive remodelling and angiogenesis in heart failure. Br J Pharmacol 166, 2348–2361 (2012); published online EpubAug (10.1111/j.1476-5381.2012.01954.x).

11. H. Kiriazis, K. Wang, Q. Xu, X. M. Gao, Z. Ming, Y. Su, X. L. Moore, G. Lambert, M. E. Gibbs, A. M. Dart, X. J. Du, Knockout of beta(1)- and beta(2)-adrenoceptors attenuates pressure overload-induced cardiac hypertrophy and fibrosis. Br J Pharmacol 153, 684–692 (2008); published online EpubFeb (10.1038/sj.bjp.0707622).

12. M. Zhao, G. Fajardo, T. Urashima, J. M. Spin, S. Poorfarahani, V. Rajagopalan, D. Huynh, A. Connolly, T. Quertermous, D. Bernstein, Cardiac pressure overload hypertrophy is differentially regulated by beta-adrenergic receptor subtypes. Am J Physiol Heart Circ Physiol 301, H1461–1470 (2011); published online EpubOct (10.1152/ajpheart.00453.2010).

13. T. M. Filtz, D. R. Grubb, T. J. McLeod-Dryden, J. Luo, E. A. Woodcock, Gq-initiated cardiomyocyte hypertrophy is mediated by phospholipase Cbeta1b. FASEB J 23, 3564–3570 (2009); published online EpubOct (10.1096/fj.09-133983).

14. D. R. Grubb, X. M. Gao, H. Kiriazis, A. Matsumoto, J. R. McMullen, X. J. Du, E. A. Woodcock, Expressing an inhibitor of PLCbeta1b sustains contractile function following pressure overload. J Mol Cell Cardiol 93, 12–17 (2016); published online EpubApr (10.1016/j.yjmcc.2016.02.015).

15. S. Malik, R. G. deRubio, M. Trembley, R. Irannejad, P. B. Wedegaertner, A. V. Smrcka, G protein betagamma subunits regulate cardiomyocyte hypertrophy through a perinuclear Golgi phosphatidylinositol 4-phosphate hydrolysis pathway. Mol Biol Cell 26, 1188–1198 (2015); published online EpubMar 15 (10.1091/mbc.E14-10-1476).

16. E. A. Oestreich, S. Malik, S. A. Goonasekera, B. C. Blaxall, G. G. Kelley, R. T. Dirksen, A. V. Smrcka, Epac and phospholipase Cepsilon regulate Ca2+ release in the heart by activation of protein kinase Cepsilon and calcium-calmodulin kinase II. J Biol Chem 284, 1514–1522 (2009); published online EpubJan 16 (10.1074/jbc.M806994200).

17. E. A. Oestreich, H. Wang, S. Malik, K. A. Kaproth-Joslin, B. C. Blaxall, G. G. Kelley, R. T. Dirksen, A. V. Smrcka, Epac-mediated activation of phospholipase C(epsilon) plays a critical role in beta-adrenergic receptor-dependent enhancement of Ca2+ mobilization in cardiac myocytes. J Biol Chem 282, 5488–5495 (2007); published online EpubFeb 23 (10.1074/jbc.M608495200).

18. L. Zhang, S. Malik, J. Pang, H. Wang, K. M. Park, D. I. Yule, B. C. Blaxall, A. V. Smrcka, Phospholipase Cepsilon hydrolyzes perinuclear phosphatidylinositol 4-phosphate to regulate cardiac hypertrophy. Cell 153, 216–227 (2013); published online EpubMar 28 (10.1016/j.cell.2013.02.047).

19. A. V. Smrcka, J. H. Brown, G. G. Holz, Role of phospholipase Cepsilon in physiological phosphoinositide signaling networks. Cell Signal 24, 1333–1343 (2012); published online EpubJun (10.1016/j.cellsig.2012.01.009).

20. G. Kadamur, E. M. Ross, Mammalian phospholipase C. Annu Rev Physiol 75, 127–154 (2013) 10.1146/annurev-physiol-030212-183750).

21. A. V. Smrcka, J. R. Hepler, K. O. Brown, P. C. Sternweis, Regulation of polyphosphoinositide-specific phospholipase C activity by purified Gq. Science 251, 804–807 (1991); published online EpubFeb 15 (10.1126/science.1846707).

22. G. G. Kelley, S. E. Reks, A. V. Smrcka, Hormonal regulation of phospholipase Cepsilon through distinct and overlapping pathways involving G12 and Ras family G-proteins. Biochem J 378, 129–139 (2004); published online EpubFeb 15 (10.1042/BJ20031370).

23. G. G. Kelley, S. E. Reks, J. M. Ondrako, A. V. Smrcka, Phospholipase C(epsilon): a novel Ras effector. EMBO J 20, 743–754 (2001); published online EpubFeb 15 (10.1093/emboj/20.4.743).

24. M. R. Wing, D. Houston, G. G. Kelley, C. J. Der, D. P. Siderovski, T. K. Harden, Activation of phospholipase C-epsilon by heterotrimeric G protein betagamma-subunits. J Biol Chem 276, 48257–48261 (2001); published online EpubDec 21 (10.1074/jbc.C100574200).

25. M. R. Wing, J. T. Snyder, J. Sondek, T. K. Harden, Direct activation of phospholipase C-epsilon by Rho. J Biol Chem 278, 41253–41258 (2003); published online EpubOct 17 (10.1074/jbc.M306904200).

26. C. A. Nash, W. Wei, R. Irannejad, A. V. Smrcka, Golgi localized beta1-adrenergic receptors stimulate Golgi PI4P hydrolysis by PLCepsilon to regulate cardiac hypertrophy. Elife 8, (2019); published online EpubAug 21 (10.7554/eLife.48167).

27. L. Zhang, S. Malik, G. G. Kelley, M. S. Kapiloff, A. V. Smrcka, Phospholipase C epsilon scaffolds to muscle-specific A kinase anchoring protein (mAKAPbeta) and integrates multiple hypertrophic stimuli in cardiac myocytes. J Biol Chem 286, 23012–23021 (2011); published online EpubJul 1 (10.1074/jbc.M111.231993).

28. J. D. Scott, L. F. Santana, A-kinase anchoring proteins: getting to the heart of the matter. Circulation 121, 1264–1271 (2010); published online EpubMar 16 (10.1161/CIRCULATIONAHA.109.896357).

29. C. A. Nash, L. M. Brown, S. Malik, X. Cheng, A. V. Smrcka, Compartmentalized cyclic nucleotides have opposing effects on regulation of hypertrophic phospholipase Cepsilon signaling in cardiac myocytes. J Mol Cell Cardiol 121, 51–59 (2018); published online EpubAug (10.1016/j.yjmcc.2018.06.002).

30. H. Duan, J. Wang, Selective transport of monoamine neurotransmitters by human plasma membrane monoamine transporter and organic cation transporter 3. J Pharmacol Exp Ther 335, 743–753 (2010); published online EpubDec (10.1124/jpet.110.170142).

31. C. D. Wright, Q. Chen, N. L. Baye, Y. Huang, C. L. Healy, S. Kasinathan, T. D. O’Connell, Nuclear alpha1-adrenergic receptors signal activated ERK localization to caveolae in adult cardiac myocytes. Circ Res 103, 992–1000 (2008); published online EpubOct 24 (10.1161/CIRCRESAHA.108.176024).

32. Y. Wang, Q. Shi, M. Li, M. Zhao, R. Reddy Gopireddy, J. P. Teoh, B. Xu, C. Zhu, K. E. Ireton, S. Srinivasan, S. Chen, P. J. Gasser, J. Bossuyt, J. W. Hell, D. M. Bers, Y. K. Xiang, Intracellular beta(1)-Adrenergic Receptors and Organic Cation Transporter 3 Mediate Phospholamban Phosphorylation to Enhance Cardiac Contractility. Circ Res 128, 246–261 (2021); published online EpubJan 22 (10.1161/CIRCRESAHA.120.317452).

33. B. Boivin, C. Lavoie, G. Vaniotis, A. Baragli, L. R. Villeneuve, N. Ethier, P. Trieu, B. G. Allen, T. E. Hebert, Functional beta-adrenergic receptor signalling on nuclear membranes in adult rat and mouse ventricular cardiomyocytes. Cardiovasc Res 71, 69–78 (2006); published online EpubJul 1 (10.1016/j.cardiores.2006.03.015).

34. G. Vaniotis, D. Del Duca, P. Trieu, C. V. Rohlicek, T. E. Hebert, B. G. Allen, Nuclear beta-adrenergic receptors modulate gene expression in adult rat heart. Cell Signal 23, 89–98 (2011); published online EpubJan (10.1016/j.cellsig.2010.08.007).

35. R. Irannejad, V. Pessino, D. Mika, B. Huang, P. B. Wedegaertner, M. Conti, M. von Zastrow, Functional selectivity of GPCR-directed drug action through location bias. Nat Chem Biol 13, 799–806 (2017); published online EpubJul (10.1038/nchembio.2389).

36. N. G. Tsvetanova, M. von Zastrow, Spatial encoding of cyclic AMP signaling specificity by GPCR endocytosis. Nat Chem Biol 10, 1061–1065 (2014); published online EpubDec (10.1038/nchembio.1665).

37. E. Devic, Y. Xiang, D. Gould, B. Kobilka, Beta-adrenergic receptor subtype-specific signaling in cardiac myocytes from beta(1) and beta(2) adrenoceptor knockout mice. Mol Pharmacol 60, 577–583 (2001); published online EpubSep (

38. Y. Xiang, E. Devic, B. Kobilka, The PDZ binding motif of the beta 1 adrenergic receptor modulates receptor trafficking and signaling in cardiac myocytes. J Biol Chem 277, 33783–33790 (2002); published online EpubSep 13 (10.1074/jbc.M204136200).

39. Y. Xiang, B. Kobilka, The PDZ-binding motif of the beta2-adrenoceptor is essential for physiologic signaling and trafficking in cardiac myocytes. Proc Natl Acad Sci U S A 100, 10776–10781 (2003); published online EpubSep 16 (10.1073/pnas.1831718100).

40. R. S. Williams, T. Bishop, Selectivity of dobutamine for adrenergic receptor subtypes: in vitro analysis by radioligand binding. J Clin Invest 67, 1703–1711 (1981); published online EpubJun (10.1172/jci110208).

41. R. P. Xiao, P. Avdonin, Y. Y. Zhou, H. Cheng, S. A. Akhter, T. Eschenhagen, R. J. Lefkowitz, W. J. Koch, E. G. Lakatta, Coupling of beta2-adrenoceptor to Gi proteins and its physiological relevance in murine cardiac myocytes. Circ Res 84, 43–52 (1999); published online EpubJan 8-22 (10.1161/01.res.84.1.43).

42. T. M. Bonacci, J. L. Mathews, C. Yuan, D. M. Lehmann, S. Malik, D. Wu, J. L. Font, J. M. Bidlack, A. V. Smrcka, Differential targeting of Gbetagamma-subunit signaling with small molecules. Science 312, 443–446 (2006); published online EpubApr 21 (10.1126/science.1120378).

43. D. M. Lehmann, A. M. Seneviratne, A. V. Smrcka, Small molecule disruption of G protein beta gamma subunit signaling inhibits neutrophil chemotaxis and inflammation. Mol Pharmacol 73, 410–418 (2008); published online EpubFeb (10.1124/mol.107.041780).

44. R. Carr, 3rd, J. Schilling, J. Song, R. L. Carter, Y. Du, S. M. Yoo, C. J. Traynham, W. J. Koch, J. Y. Cheung, D. G. Tilley, J. L. Benovic, beta-arrestin-biased signaling through the beta2-adrenergic receptor promotes cardiomyocyte contraction. Proc Natl Acad Sci U S A 113, E4107-4116 (2016); published online EpubJul 12 (10.1073/pnas.1606267113).

45. S. Maudsley, K. L. Pierce, A. M. Zamah, W. E. Miller, S. Ahn, Y. Daaka, R. J. Lefkowitz, L. M. Luttrell, The beta(2)-adrenergic receptor mediates extracellular signal-regulated kinase activation via assembly of a multi-receptor complex with the epidermal growth factor receptor. J Biol Chem 275, 9572–9580 (2000); published online EpubMar 31 (10.1074/jbc.275.13.9572).

46. I. M. Kim, D. G. Tilley, J. Chen, N. C. Salazar, E. J. Whalen, J. D. Violin, H. A. Rockman, Beta-blockers alprenolol and carvedilol stimulate beta-arrestin-mediated EGFR transactivation. Proc Natl Acad Sci U S A 105, 14555–14560 (2008); published online EpubSep 23 (10.1073/pnas.0804745105).

47. L. A. Grisanti, J. A. Talarico, R. L. Carter, J. E. Yu, A. A. Repas, S. W. Radcliffe, H. A. Tang, C. A. Makarewich, S. R. Houser, D. G. Tilley, beta-Adrenergic receptor-mediated transactivation of epidermal growth factor receptor decreases cardiomyocyte apoptosis through differential subcellular activation of ERK1/2 and Akt. J Mol Cell Cardiol 72, 39–51 (2014); published online EpubJul (10.1016/j.yjmcc.2014.02.009).

48. Y. Kwon, S. Mehta, M. Clark, G. Walters, Y. Zhong, H. N. Lee, R. K. Sunahara, J. Zhang, Non-canonical beta-adrenergic activation of ERK at endosomes. Nature 611, 173–179 (2022); published online EpubNov (10.1038/s41586-022-05343-3).

49. L. E. Gimenez, F. Baameur, S. J. Vayttaden, R. B. Clark, Salmeterol Efficacy and Bias in the Activation and Kinase-Mediated Desensitization of beta2-Adrenergic Receptors. Mol Pharmacol 87, 954–964 (2015); published online EpubJun (10.1124/mol.114.096800).

50. R. H. Moore, E. E. Millman, V. Godines, N. A. Hanania, T. M. Tran, H. Peng, B. F. Dickey, B. J. Knoll, R. B. Clark, Salmeterol stimulation dissociates beta2-adrenergic receptor phosphorylation and internalization. Am J Respir Cell Mol Biol 36, 254–261 (2007); published online EpubFeb (10.1165/rcmb.2006-0158OC).

51. V. Patki, D. C. Lawe, S. Corvera, J. V. Virbasius, A. Chawla, A functional PtdIns(3)P-binding motif. Nature 394, 433–434 (1998); published online EpubJul 30 (10.1038/28771).

52. M. T. Kunkel, A. Toker, R. Y. Tsien, A. C. Newton, Calcium-dependent regulation of protein kinase D revealed by a genetically encoded kinase activity reporter. J Biol Chem 282, 6733–6742 (2007); published online EpubMar 2 (10.1074/jbc.M608086200).

53. N. Frey, E. N. Olson, Cardiac hypertrophy: the good, the bad, and the ugly. Annu Rev Physiol 65, 45–79 (2003) 10.1146/annurev.physiol.65.092101.142243).

54. M. Bohm, T. Eschenhagen, P. Gierschik, K. Larisch, H. Lensche, U. Mende, W. Schmitz, P. Schnabel, H. Scholz, M. Steinfath, et al., Radioimmunochemical quantification of Gi alpha in right and left ventricles from patients with ischaemic and dilated cardiomyopathy and predominant left ventricular failure. J Mol Cell Cardiol 26, 133–149 (1994); published online EpubFeb (10.1006/jmcc.1994.1017).

55. A. M. Feldman, A. E. Cates, W. B. Veazey, R. E. Hershberger, M. R. Bristow, K. L. Baughman, W. A. Baumgartner, C. Van Dop, Increase of the 40,000-mol wt pertussis toxin substrate (G protein) in the failing human heart. J Clin Invest 82, 189–197 (1988); published online EpubJul (10.1172/JCI113569).

56. K. Lorenz, J. P. Schmitt, E. M. Schmitteckert, M. J. Lohse, A new type of ERK1/2 autophosphorylation causes cardiac hypertrophy. Nat Med 15, 75–83 (2009); published online EpubJan (10.1038/nm.1893).

57. M. Mutlak, M. Schlesinger-Laufer, T. Haas, R. Shofti, N. Ballan, Y. E. Lewis, M. Zuler, Y. Zohar, L. H. Caspi, I. Kehat, Extracellular signal-regulated kinase (ERK) activation preserves cardiac function in pressure overload induced hypertrophy. Int J Cardiol 270, 204–213 (2018); published online EpubNov 1 (10.1016/j.ijcard.2018.05.068).

58. N. H. Purcell, B. J. Wilkins, A. York, M. K. Saba-El-Leil, S. Meloche, J. Robbins, J. D. Molkentin, Genetic inhibition of cardiac ERK1/2 promotes stress-induced apoptosis and heart failure but has no effect on hypertrophy in vivo. Proc Natl Acad Sci U S A 104, 14074–14079 (2007); published online EpubAug 28 (10.1073/pnas.0610906104).

59. I. Kehat, J. Davis, M. Tiburcy, F. Accornero, M. K. Saba-El-Leil, M. Maillet, A. J. York, J. N. Lorenz, W. H. Zimmermann, S. Meloche, J. D. Molkentin, Extracellular signal-regulated kinases 1 and 2 regulate the balance between eccentric and concentric cardiac growth. Circ Res 108, 176–183 (2011); published online EpubJan 21 (10.1161/CIRCRESAHA.110.231514).

60. M. Silberbach, T. Gorenc, R. E. Hershberger, P. J. Stork, P. S. Steyger, C. T. Roberts, Jr., Extracellular signal-regulated protein kinase activation is required for the anti-hypertrophic effect of atrial natriuretic factor in neonatal rat ventricular myocytes. J Biol Chem 274, 24858–24864 (1999); published online EpubAug 27 (10.1074/jbc.274.35.24858).

61. H. Lavoie, J. Gagnon, M. Therrien, ERK signalling: a master regulator of cell behaviour, life and fate. Nat Rev Mol Cell Biol 21, 607–632 (2020); published online EpubOct (10.1038/s41580-020-0255-7).

62. K. Eichel, M. von Zastrow, Subcellular Organization of GPCR Signaling. Trends Pharmacol Sci 39, 200–208 (2018); published online EpubFeb (10.1016/j.tips.2017.11.009).

63. S. Ahn, S. K. Shenoy, H. Wei, R. J. Lefkowitz, Differential kinetic and spatial patterns of beta-arrestin and G protein-mediated ERK activation by the angiotensin II receptor. J Biol Chem 279, 35518–35525 (2004); published online EpubAug 20 (10.1074/jbc.M405878200).

64. Z. G. Goldsmith, D. N. Dhanasekaran, G protein regulation of MAPK networks. Oncogene 26, 3122–3142 (2007); published online EpubMay 14 (10.1038/sj.onc.1210407).

65. S. Rajagopal, K. Rajagopal, R. J. Lefkowitz, Teaching old receptors new tricks: biasing seven-transmembrane receptors. Nat Rev Drug Discov 9, 373–386 (2010); published online EpubMay (10.1038/nrd3024).

66. A. K. Shukla, J. D. Violin, E. J. Whalen, D. Gesty-Palmer, S. K. Shenoy, R. J. Lefkowitz, Distinct conformational changes in beta-arrestin report biased agonism at seven-transmembrane receptors. Proc Natl Acad Sci U S A 105, 9988–9993 (2008); published online EpubJul 22 (10.1073/pnas.0804246105).

67. J. Fielitz, M. S. Kim, J. M. Shelton, X. Qi, J. A. Hill, J. A. Richardson, R. Bassel-Duby, E. N. Olson, Requirement of protein kinase D1 for pathological cardiac remodeling. Proc Natl Acad Sci U S A 105, 3059–3063 (2008); published online EpubFeb 26 (10.1073/pnas.0712265105).

68. A. Roy, J. Ye, F. Deng, Q. J. Wang, Protein kinase D signaling in cancer: A friend or foe? Biochim Biophys Acta Rev Cancer 1868, 283–294 (2017); published online EpubAug (10.1016/j.bbcan.2017.05.008).

69. A. Chesley, M. S. Lundberg, T. Asai, R. P. Xiao, S. Ohtani, E. G. Lakatta, M. T. Crow, The beta(2)-adrenergic receptor delivers an antiapoptotic signal to cardiac myocytes through G(i)-dependent coupling to phosphatidylinositol 3’-kinase. Circ Res 87, 1172–1179 (2000); published online EpubDec 8 (10.1161/01.res.87.12.1172).

70. H. Chen, N. Ma, J. Xia, J. Liu, Z. Xu, beta2-Adrenergic receptor-induced transactivation of epidermal growth factor receptor and platelet-derived growth factor receptor via Src kinase promotes rat cardiomyocyte survival. Cell Biol Int 36, 237–244 (2012); published online EpubMar 1 (10.1042/CBI20110162).

71. P. A. Poole-Wilson, K. Swedberg, J. G. Cleland, A. Di Lenarda, P. Hanrath, M. Komajda, J. Lubsen, B. Lutiger, M. Metra, W. J. Remme, C. Torp-Pedersen, A. Scherhag, A. Skene, I. Carvedilol Or Metoprolol European Trial, Comparison of carvedilol and metoprolol on clinical outcomes in patients with chronic heart failure in the Carvedilol Or Metoprolol European Trial (COMET): randomised controlled trial. Lancet 362, 7–13 (2003); published online EpubJul 5 (10.1016/S0140-6736(03)13800-7).

72. J. W. Wisler, S. M. DeWire, E. J. Whalen, J. D. Violin, M. T. Drake, S. Ahn, S. K. Shenoy, R. J. Lefkowitz, A unique mechanism of beta-blocker action: carvedilol stimulates beta-arrestin signaling. Proc Natl Acad Sci U S A 104, 16657–16662 (2007); published online EpubOct 16 (10.1073/pnas.0707936104).

73. I. Ahmet, M. Krawczyk, W. Zhu, A. Y. Woo, C. Morrell, S. Poosala, R. P. Xiao, E. G. Lakatta, M. I. Talan, Cardioprotective and survival benefits of long-term combined therapy with beta2 adrenoreceptor (AR) agonist and beta1 AR blocker in dilated cardiomyopathy postmyocardial infarction. J Pharmacol Exp Ther 325, 491–499 (2008); published online EpubMay (10.1124/jpet.107.135335).

74. M. I. Talan, I. Ahmet, R. P. Xiao, E. G. Lakatta, beta(2) AR agonists in treatment of chronic heart failure: long path to translation. J Mol Cell Cardiol 51, 529–533 (2011); published online EpubOct (10.1016/j.yjmcc.2010.09.019).

75. H. Stenmark, R. Aasland, B. H. Toh, A. D’Arrigo, Endosomal localization of the autoantigen EEA1 is mediated by a zinc-binding FYVE finger. J Biol Chem 271, 24048–24054 (1996); published online EpubSep 27 (10.1074/jbc.271.39.24048).

76. J. Liu, T. E. Hughes, W. C. Sessa, The first 35 amino acids and fatty acylation sites determine the molecular targeting of endothelial nitric oxide synthase into the Golgi region of cells: a green fluorescent protein study. J Cell Biol 137, 1525–1535 (1997); published online EpubJun 30 (10.1083/jcb.137.7.1525).

77. D. S. Auerbach, J. Jones, B. C. Clawson, J. Offord, G. M. Lenk, I. Ogiwara, K. Yamakawa, M. H. Meisler, J. M. Parent, L. L. Isom, Altered cardiac electrophysiology and SUDEP in a model of Dravet syndrome. PLoS One 8, e77843 (2013)10.1371/journal.pone.0077843).

